# A cell-free system for mitotically-stimulated TMEJ

**DOI:** 10.64898/2026.10.26.762058

**Authors:** Yufan Wu, Martha J. Longley, Geylani Can, Johannes C. Walter, Joseph J. Loparo

**Affiliations:** Department of Biological Chemistry and Molecular Pharmacology, Blavatnik Institute, Harvard Medical School, Boston, MA 02115, USA; Howard Hughes Medical Institute

**Author notes:** These authors contributed equally (Johannes C. Walter, Joseph J. Loparo).

## Abstract

Double-stranded DNA breaks (DSBs) are cytotoxic and must be repaired to avoid cell death or mutagenesis. DSBs that form in the S or G2 phases of the cell cycle undergo end resection, creating substrates for homologous recombination (HR) and alternative end joining pathways. Theta-mediated end joining (TMEJ) is a DNA polymerase θ (Pol θ)-dependent repair pathway that joins resected DSB ends by annealing short regions of microhomology. TMEJ becomes particularly important when HR is compromised or during mitosis. Despite intensive investigation, the mechanism of TMEJ remains poorly understood, in part due to the lack of an efficient cell-free system that recapitulates this reaction. Here, we address this deficit using *Xenopus* egg extracts. We show that a linear DNA template containing a 3^’^ single-stranded DNA overhang with terminal microhomology undergoes efficient intermolecular joining in nucleoplasmic extracts derived from *Xenopus* eggs. Joining is abolished upon Pol θ depletion, and restoration of joining requires Pol θ that is proficient in both ATPase and DNA polymerase activities. Furthermore, joining is accelerated by recombinant Cyclin B–CDK1, which also enhances the recruitment of Pol θ to single-stranded DNA ends. This assay provides a biochemically tractable approach for dissecting the mechanism and regulation of TMEJ in a physiologically complex environment.

## INTRODUCTION

DNA double-strand breaks (DSBs) are deleterious lesions whose repair prevents cell death, genomic instability, and tumorigenesis [1]. In addition to classical non-homologous end joining (c-NHEJ) and homologous recombination (HR), metazoan cells employ theta-mediated end joining (TMEJ), which uses DNA polymerase theta (Pol θ) to join resected DNA ends by annealing short regions of microhomology. Although initially viewed as a backup pathway, TMEJ is increasingly recognized as an important contributor to genome maintenance [2–8]. TMEJ contributes to the formation and maintenance of extrachromosomal DNA, a driver of oncogene amplification and therapy resistance [9]. Moreover, subtypes of breast and ovarian cancers with HR defects depend on TMEJ for survival [10–12]. Inhibition of Pol θ sensitizes cancer cells to ionizing radiation [13] and other damaging agents [12,14], making it an attractive target for cancer therapy.

TMEJ acts on resected DSB ends generated by the MRE11-RAD50-NBS1 (MRN) complex and CtIP [15–17]. Pol θ then uses exposed microhomologies within the 3^’^ ssDNA overhangs to coordinate end synapsis and repair synthesis [18–20]. Pol θ contains an N-terminal helicase-like domain and a C-terminal A-family polymerase domain connected by a disordered central domain [21,22]. Its helicase-like domain possesses ATPase activity that counteracts ssDNA-bound RPA and promotes end capture and juxtaposition [23,24], whereas the polymerase domain stabilizes mismatched or weakly annealed microhomologies and initiates DNA synthesis [25,26]. Nucleases then remove non-homologous DNA tails [27,28], followed by further strand extension [29,30] and ligation to complete end joining [31]. As a result, TMEJ produces characteristic repair junctions enriched in microhomology-associated deletions and templated insertions [19,32].

The requirement for resected DNA ends places TMEJ under strong cell-cycle control. As DNA end resection is favored when cyclin-dependent kinase activity is high, TMEJ has traditionally been associated with S and G2 phases [15], where it contributes to the repair of replication-associated lesions [5,20,33–37]. More recent work has revealed a particularly prominent role for TMEJ as cells approach and enter mitosis [38–43]. This mitotic activation has been linked directly to Pol θ recruitment [41,42]. PLK1-dependent phosphorylation of the Pol θ central domain promotes its interaction with TOPBP1 and accumulation at mitotic DSBs [42]. Meanwhile, the mitosis-specific factor RHINO independently interacts with Pol θ and promotes its efficient accumulation at mitotic DSBs [41]. Together, these studies support a model in which TMEJ becomes particularly important for resolving DSBs that persist into mitosis, when HR and c-NHEJ are suppressed. However, it is unclear to what extent the apparent cell-cycle dependence of TMEJ reflects direct regulation of TMEJ itself, rather than changes in substrate availability and competing repair pathways. More broadly, how the distinct activities of Pol θ and its interactions with other repair factors are regulated to execute TMEJ in cells remains incompletely understood.

Biochemical approaches have shed light on the mechanism of TMEJ. Early studies using the isolated helicase-like and polymerase domains of Pol θ identified structural features underlying their biochemical activities and defined key substrate requirements for TMEJ [23,25,44–52]. Subsequent work with purified full-length Pol θ revealed that tethering the helicase-like and polymerase domains promotes DNA synapsis and efficient TMEJ of ssDNA overhangs resembling resected DSBs [53]. More recently, a reconstituted system comprising RPA, Pol δ, and a Pol θ variant with a shortened central linker recapitulated key early steps of TMEJ [54]. Furthermore, to capture the broader biochemical environment in which TMEJ operates, *Xenopus* egg extracts were used to characterize Pol θ-dependent repair of 5′ DNA-protein crosslinks [55] and joining of DNA ends generated upon mitotic replication fork collapse [43]. However, in both settings, joining activity was low and coupled to upstream lesion processing or replication-associated fork breakage, making it difficult to examine the regulation of the core joining reaction itself.

Here, we established a cell-free TMEJ system in *Xenopus* egg extracts using a defined, pre-resected linear DNA substrate containing a 3′ ssDNA overhang. End joining was Pol θ dependent and required both its ATPase and polymerase activities. Joining was already robust in interphase extracts, and addition of recombinant Cyclin B–CDK1 to create a mitotic environment accelerated it approximately two-fold. This acceleration was accompanied by Pol θ phosphorylation and by increased recruitment of Pol θ to single-stranded DNA, suggesting that mitotic kinase activity enhances TMEJ at least in part by promoting Pol θ recruitment. Together, our findings establish *Xenopus* egg extracts as a tractable system for dissecting the mechanism and regulation of TMEJ.

## RESULTS

### Efficient Pol θ-mediated end joining in frog egg extracts

Previous work showed that depletion of Pol θ from whole *Xenopus* egg lysates abolished the joining of linear DNA molecules containing 5′ DNA–protein crosslinks, but re-addition of recombinant Pol θ purified from yeast restored only limited joining activity [55]. To establish a more robust, Pol θ-dependent cell-free system for TMEJ, we explored the use of concentrated nucleoplasmic extract (NPE) from unfertilized *Xenopus* eggs, since it supports many other forms of DNA repair [56–60]. We first examined how different DNA end structures influenced DSB repair pathway choice in NPE. Blunt-ended linear DNA was largely degraded, consistent with extensive 5^’^ end resection outcompeting end joining (Fig. S1A, lanes 1-4). Mre11 depletion stabilized this substrate and promoted formation of mostly supercoiled products, which disappeared upon DNAPK inhibition, indicative of NHEJ (Fig. S1A, lanes 5-12) [61]. We next used a linear DNA substrate containing 3^’^ ssDNA overhangs with terminal microhomology to mimic a resected DSB intermediate. Because we wished to preserve the defined end structure of this substrate, we depleted Mre11 to limit further end resection. In Mre11-depeted NPE, this substrate generated predominantly higher molecular-weight joining products that were greatly reduced by Pol θ depletion, consistent with TMEJ (Fig. S1B, lanes 5-12). To simplify the end-joining products, we took advantage of the observation that hairpin-capped DNA ends block joining in Mre11-depleted extract (Fig. S1C). We therefore generated a substrate with one microhomology-containing ssDNA end and one hairpin-capped end (Fig. 1A). This design restricted productive intermolecular joining to a single event and yielded a discrete, dimeric product (Fig. 1B, lanes 1-4). Dimer formation was abolished by Pol θ depletion but was unaffected by depletion of the core NHEJ factor Ku (Fig. 1B, lanes 5-12, and Fig. 1C).

**Figure 1.**
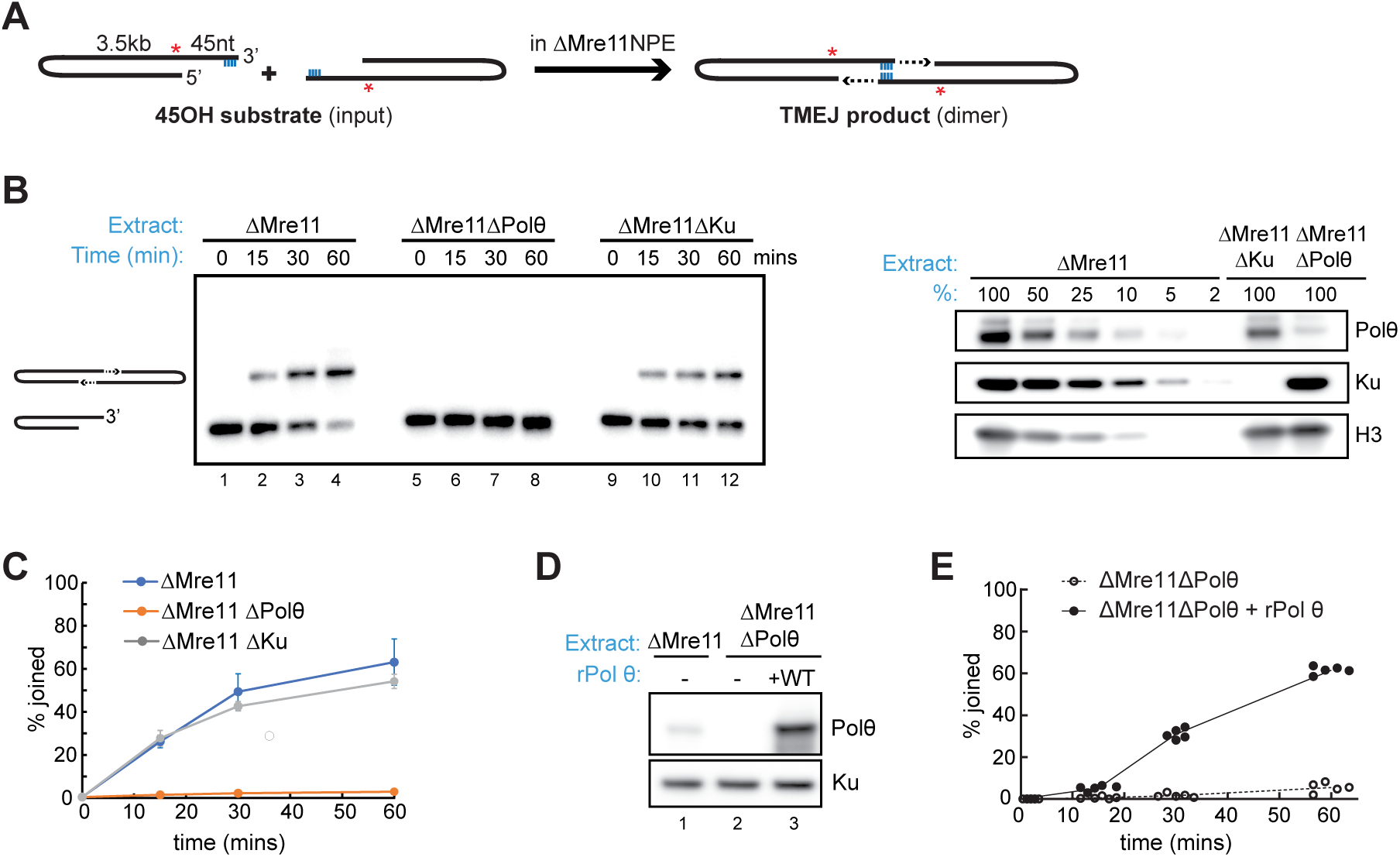
Robust Pol θ-mediated end joining in *Xenopus* egg extract. **(A)** Diagram of the TMEJ assay using the “45OH substrate”. Red star, radiolabel. Blue line, terminal microhomology (CCGG-3’). **(B)** Left: 45OH substrate was incubated with the indicated NPE extracts (optionally supplemented with 10 μM DNA-PK inhibitor NU-7441), separated on a native agarose gel, and subjected to autoradiography. Right: The indicated extracts were blotted with the indicated antibodies. **(C)** Quantification of joining efficiency of the experiment shown in (B). N=3. Mean ± SEM. **(D)** The indicated extracts were blotted with the indicated antibodies. Depleted NPE was supplemented with wild-type Pol θ expressed using a tobacco-cell transcription-translation extract (TTE). **(E)** Quantification of 45OH joining efficiency in the indicated extracts supplemented with Pol θ expressed in one lot (#1) of tobacco-cell extract. N=5 experiments.

To confirm that the joining defect in Pol θ-depleted NPE was specific, we attempted to restore end joining with recombinant Pol θ. *Xenopus* Pol θ produced in insect cells or human Pol θ expressed in human cells did not restore joining (data not shown), likely because active full-length Pol θ could not be obtained at sufficiently high concentration [54,55]. We therefore expressed Pol θ in a Tobacco-cell transcription-translation extract (TTE) [62], allowing us to add the protein to extracts at or exceeding endogenous levels (Fig. 1D, compare lanes 1 and 3). This TTE-expressed Pol θ restored robust dimer formation in Pol θ-depleted NPE to levels seen in mock-depleted extract, demonstrating specificity of the depletion (compare Fig. 1E with Fig. 1C). Although rescue efficiency varied depending on the batch of TTE used to express Pol θ, wild-type Pol θ consistently restored DNA joining to levels above background (Fig. 1E and Fig. S2). Collectively, these results establish an extract-based assay in which Pol θ-mediated end joining generates a discrete dimeric product from a defined pre-resected DNA substrate.

### Cell-free TMEJ requires the ATPase and polymerase activities of Pol θ

Previous biochemical and cellular studies have established an essential role for Pol θ polymerase activity in TMEJ [10,20,26,33]. In contrast, the requirement for ATPase activity of the helicase-like domain depends on the cellular context [5,10,20,24]. To define the contribution of these enzymatic activities to cell-free TMEJ, we first treated extracts with ART558, an allosteric inhibitor of the Pol θ polymerase domain [63]. ART558 strongly inhibited dimer formation of the TMEJ substrate but had little effect on NHEJ of a blunt-ended DNA substrate (Fig. 2A and Fig. S3), indicating a requirement for Pol θ polymerase activity. Consistent with this result, a recombinant Pol θ variant carrying inactivating mutations in the conserved polymerase domain (DY2286/7AA) [64] failed to restore joining in Pol θ-depleted extracts (Fig. 2B), despite being added back at concentrations equivalent to the WT protein (Fig. 2C, compare lanes 3 and 4).

**Figure 2.**
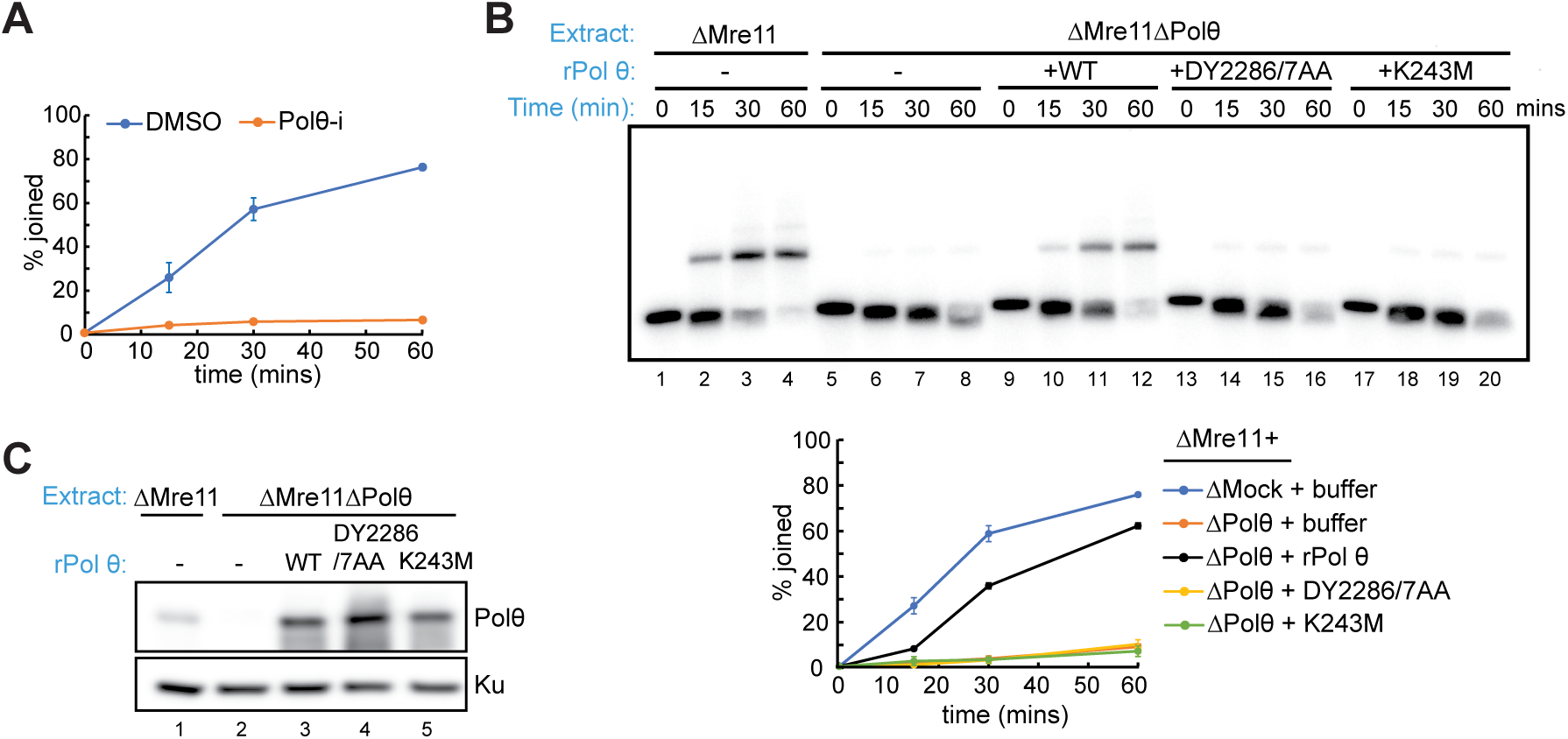
Pol θ-mediated end joining in *Xenopus* egg extracts requires both the ATPase and polymerase activities of Pol θ. **(A)** Quantification of 45OH joining efficiency in Mre11-depleted NPE in the presence of DMSO or 10 µM Pol θ inhibitor ART558. N=3 experiments. Mean ± SEM. Representative gel shown in Fig. S3. **(B)** Top: Joining of the 45OH substrate in the indicated extracts supplemented with the indicated Pol θ constructs. Bottom: Quantification of joining efficiency, N=3 experiments. Mean ± SEM. **(C)** The extracts used in (B) were blotted with the indicated antibodies.

To examine the requirement for the helicase-like domain, we introduced a mutation into the conserved ATP-binding site of the Walker A motif (K243M), which abolishes ATPase activity [20]. In contrast to wild-type Pol θ, this Pol θ mutant also failed to restore dimer formation in Pol θ-depleted extracts (Fig. 2B and Fig. 2C). Thus, although ATPase activity can be partially dispensable for TMEJ in certain cellular contexts, it is required for efficient TMEJ in NPE. Together, these results demonstrate that both the polymerase and ATPase activities of Pol θ are required for efficient cell-free TMEJ.

### Cyclin B–CDK1 enhances cell-free TMEJ

Recent studies suggest that TMEJ is particularly important during mitosis, when HR and NHEJ are inactive. To determine how mitotic kinase activity affects TMEJ, we preincubated NPE with recombinant Cyclin B– CDK1 (B–CDK1) [43,65] before addition of the DNA substrate. B–CDK1 accelerated joining approximately two-fold at early time points, although the extent of joining at 60 min was similar across conditions (Fig. 3A and Fig. S4). In this setting, we observed little variation in the efficiency with which Pol θ-depletion was rescued using different batches of TTE-expressed Pol θ, including the same preparations of Pol θ that showed variable rescue in the interphase setting (Fig. S5).

**Figure 3.**
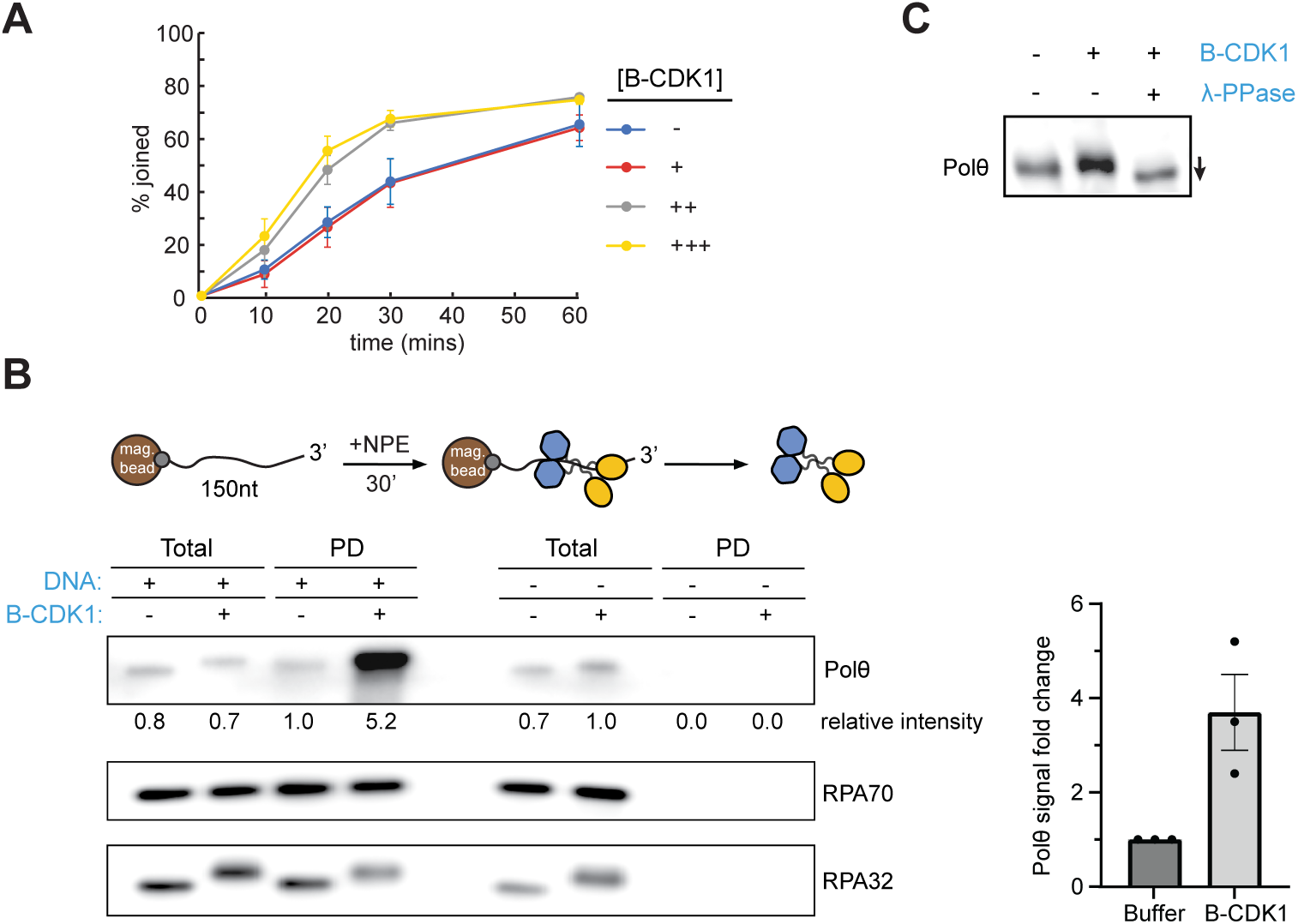
Cyclin B–CDK1 accelerates TMEJ and increases Pol θ recruitment to ssDNA. **(A)** Quantification of joining efficiency in Mre11-depleted NPE supplemented with increasing amounts of Cyclin B–CDK1 (0, 3, 8.3 and 25 ng/μl). N=3 experiments. Mean ± SEM. A representative gel is shown in Fig. S4. **(B)** Left: ssDNA pull-down assay. A 5′ biotinylated 150 nt ssDNA with a free 3′ end was immobilized on magnetic beads and incubated with NPE supplemented with buffer or Cyclin B–CDK1 as indicated. The beads were subsequently washed and proteins were eluted for Western blot analysis. A representative Western blot of whole extract (total) and proteins recovered from ssDNA (PD) is shown. N=3 experiments. Magnetic beads without ssDNA attached (no DNA) served as controls for nonspecific binding signals. Right: Quantification of the ssDNA pull-down assay in NPE supplemented with buffer or Cyclin B–CDK1. N=3 experiments. Mean ± SEM. **(C)** NPE was supplemented with buffer or Cyclin B–CDK1, with the latter additionally treated with λ phosphatase where indicated, and analyzed by Phos-tag gel electrophoresis and immunoblotting for Pol θ.

To determine whether stimulation by B–CDK1 was associated with enhanced recruitment of Pol θ to resected DNA ends, we immobilized a 150 nt single strand DNA (ssDNA) with a free 3′ end to streptavidin magnetic beads, incubated the substrate in egg extracts, and analyzed DNA-associated proteins after recovery (Fig. 3B). RPA association with ssDNA was unchanged by B–CDK1 treatment. In contrast, we observed a 3.7-fold increase in the recruitment of Pol θ to the DNA in the presence of B–CDK1. In addition, B–CDK1 induced a noticeable mobility shift of Pol θ that was abolished by phosphatase (Fig. 3C), consistent with phosphorylation of Pol θ in B–CDK1 treated extracts. Together, these results show that B–CDK1 enhances cell-free TMEJ and raise the possibility that increased recruitment of Pol θ to DNA may contribute to this enhancement.

## DISCUSSION

Here, we report that *Xenopus* egg extracts support robust TMEJ, with over 50% joining in an hour. Both the ATPase and DNA polymerase activities of Pol θ are required for TMEJ in this system. TMEJ occurs in a complex, quasi-cellular environment containing physiological protein concentrations where the contribution of any protein can be determined via immunodepletion. B–CDK1 treatment enhances TMEJ approximately two-fold and is accompanied by increased Pol θ phosphorylation and recruitment to the DNA substrate. These findings establish a physiologically complex reconstitution of TMEJ that will lend new insights into the mechanism of TMEJ. Indeed, we recently used this approach to show that RPA is essential for TMEJ [66].

Our findings support existing evidence that the central events in TMEJ require not only Pol θ’s polymerization activity, but also its ATPase activity. While studies from minimally reconstituted systems have demonstrated that Pol θ’s helicase-like domain counteracts ssDNA-bound RPA and facilitates strand annealing [23,24], whether this enzymatic activity is essential in cells remains debated. ATPase-deficient Pol θ rescued bleomycin sensitivity in mouse embryonic fibroblasts (MEFs) [20]. It also largely preserved TMEJ activity on extrachromosomal substrates in MEFs [10] and at ionizing radiation-induced DSBs in Drosophila [5], although internal microhomology usage was reduced. In contrast, ATPase activity was required for interstrand crosslink repair in Drosophila [5] and contributed to the proliferation of HR-deficient mouse embryonic stem cells and chromosomal translocation formation [24]. These findings indicate that the requirement for Pol θ ATPase activity is highly context dependent in cellular systems. Our cell-free TMEJ assay uses a pre-resected substrate containing a 4-nt terminal microhomology, thereby minimizing the need for end resection and internal microhomology search, providing a more direct readout of the core TMEJ reaction. Under these conditions, ATPase-dead Pol θ failed to support joining, despite the presence of a terminal microhomology. This finding suggests there is an intrinsic requirement for ATP hydrolysis in TMEJ, potentially to fully remove ssDNA-bound RPA and facilitate annealing [23,24].

Our findings also provide mechanistic insight into the apparent cell-cycle regulation of TMEJ. Robust joining was observed in interphase extract, and it was further accelerated upon addition of B–CDK1, consistent with recent evidence that TMEJ becomes prominent during mitosis. Importantly, the substantial activity observed in interphase extract indicates that the core enzymatic machinery of TMEJ does not require mitotic activation. In line with this, Pol θ-mediated repair of DSBs in B lymphocytes lacking NHEJ coincides with the G1-to-S phase transition [2], supporting the idea that the pathway is competent outside mitosis when suitable substrates are available. Following addition of B–CDK1, the recruitment of Pol θ to ssDNA was markedly increased, suggesting that the mitotic enhancement arises from increased Pol θ engagement with DNA.

Our findings show that mitotic conditions enhance Pol θ association with resected DNA ends relative to interphase. It will be interesting to determine whether this enhancement is mediated via TOPBP1, RHINO and CIP2A [41,67,68], or whether other mechanisms are at play. Beyond mitotic regulation, ATR kinase activity has recently been implicated in TMEJ during R2 retrotransposon integration [65], raising the question of whether DNA damage response kinases also modulate Pol θ recruitment, independently of or in concert with mitotic kinase activity. Our cell-free system provides a tractable platform for dissecting these kinase-dependent mechanisms.

## ACKNOWLEDGEMENTS

This work was supported by Harvard Medical School Dean’s innovation grant (to J.J.L and J.C.W) and National Institutes of Health grant R01CA272436 (to J.J.L). J.C.W. was supported by NIH grant HL098316. J.C.W. is an American Cancer Society research professor and a member of the Howard Hughes Medical Institute. Y.W was supported by BCMP-Merck fellowship. We thank members of the Loparo and Walter lab for useful discussions and critical comments on the paper.

## METHODS

### Chemicals & Reagents

ART558 was purchased from Tocris Bioscience. Cyclin B– CDK1 was purchased from EMD Millipore (Cat# 14-450M). Oligonucleotides were purchased from Integrated DNA Technologies.

### Preparation of DNA substrates

To make the TMEJ substrate, a 5.7 kb plasmid (a derivative of pACE) was linearized with NotI and BamHI (New England Biolabs, NEB). The resulting 3.5 kb fragment was separated from the 2.2kb fragment on a 0.8% 1X TBE agarose gel and recovered by electroelution and ethanol precipitation. The overhang and hairpin adapters were ligated to each side of this backbone fragment to generate the described DNA substrates. Oligonucleotide stocks (10 μM) were first phosphorylated with T4 PNK (NEB). To generate the radioactive overhang adaptor, oligonucleotide (1) was first phosphorylated for 30 mins with γ-32P-ATP (Perkin Elmer), and subsequently 50 μM cold ATP was added and the reaction was allowed to proceed a further 15 mins to ensure maximal phosphorylation. To generate the hairpin adaptor, the phosphorylated oligonucleotide (2) was diluted in half in annealing buffer (NEBuffer™ 2), heated at 95°C for 5 mins then directly placed on ice to promote self-annealing for at least 20 mins. The hairpin (250 nM) and overhang adapters (375 nM cold and 125 nM radiolabeled) were ligated to the backbone fragment (25 nM) with T4 DNA ligase at 16°C for 5 hours. The desired 3.5 kb product was separated on a 0.8% 1x TBE agarose gel and recovered with the QIAquick gel extraction kit (QIAGEN). Blunt end and hairpin end substrates (used in Fig. S1) were prepared using previous protocols with small modifications [61,69].

### Extract and ensemble joining assay

Experiments involving adult female (Nasco Cat# LM0053MX) and male (Nasco Cat# LM00715MX) *Xenopus laevis* were approved by the Harvard Medical Area Institutional Animal Care and Use Committee (IACUC) and conform to all relevant regulatory standards. All frogs were healthy, maintained at 18°C on a 12-h light cycle and a biweekly feeding schedule. The water was purified by reverse osmosis and conditioned with rock salt approved for aquaculture. Female frogs were handled and ovulated according to approved protocols. Male frogs were not involved in previous procedures.

*Xenopus* egg extracts were prepared as described [70,71]. The ensemble gel-based end joining time course was performed similarly to a previous protocol [61,72]. For each reaction condition, extract was supplemented with a 30x ATP regeneration mixture (65 mM ATP, 650 mM phosphocreatine, 160 ng/μL creatine phosphokinase) to a 1x final concentration and 10-30 ng/μL closed circular ‘carrier’ DNA that is required for joining of dilute linear substrates as well as for DNA replication in extract [61,73]. Joining reactions were initiated by addition of 3 ng/μL radiolabeled linear DNA substrate and carried out at room temperature for the indicated time. Time points were taken by adding 2 μL of the reaction mixture to 5 μL of stop solution (80 mM Tris, pH 8.0, 8 mM EDTA, 0.13% phosphoric acid, 10% Ficoll, 5% SDS, and 0.2% bromophenol blue). The 0 min time point was taken and mixed with stop solution immediately after adding the radiolabeled substrate to the reaction and mixing. All reaction samples were digested for 1-2 hr at 37°C with 1.4 mg/mL proteinase K. Digested samples were then run on a Tris-borate-EDTA 0.8% agarose gel. The gel was pressed and dried onto a HyBond-XL nylon membrane (GE Healthcare) and exposed to a storage phosphor screen. Exposed screens were scanned using a Typhoon FLA 7000 imager (GE Healthcare).

For joining reactions in the presence of Cyclin B– CDK1 (B–CDK1), extracts were supplemented with the indicated amounts of B–CDK1 (0, 3, 8.3, or 25 ng/μl final concentration for the titration experiment in Fig. 3A; and incubated at room temperature for 15 min. The DNA substrate was then added to initiate the reaction.

### Pol θ Rescue experiments

For Pol θ rescue experiments, *Xenopus laevis* Pol θ was cloned into the pALiCE01 backbone for expression in ALiCE® eukaryotic cell-free protein expression system (LenioBio). Protein expression conditions were based on the manufacturer’s instructions with minor modifications. Proteins were expressed in the ALiCE cell-free system at a plasmid concentration of 7 nM for 44-48 hr at 25°C and 700 rpm in an Eppendorf ThermoMixer, in a humidified environment (relatively humidity >70%). The samples were centrifuged at 16,000 x g at 4°C for 5 min following expression, and the supernatant was aliquoted, flash-frozen and kept at −80°C until use. Each ALiCE-expressed Pol θ variant was mixed with egg extract at a 1:10 (v/v) ratio and incubated at room temperature for 15 min. The DNA substrate was subsequently added to initiate the reaction. ALiCE without an expression vector was added back to conditions where protein was not added to ensure that the volume and composition of each reaction were directly comparable.

For rescue experiments in the presence of B–CDK1, ALiCE-expressed Pol θ was mixed with egg extract supplemented with 16 ng/μl B–CDK1. The mixture was incubated at room temperature for 15 min, followed by the addition of the DNA substrate.

### Immunodepletion and antibody

Immunodepletion using antibodies against *Xenopus laevis* Mre11 (see below), Pol θ (antigen: residues 1212 to 1506, Abgent) [74] and Ku [61] were performed as described. Briefly, Protein A Sepharose Fast Flow beads (GE Healthcare) were incubated with antibodies at 4°C overnight. Five volumes of pre-cleared egg extract were then mixed with one volume of antibody-bound Sepharose beads. For Ku, Mre11 and Pol θ depletion of egg extract, three rounds of depletion were performed at 4°C for 1 hr each. For each round, a mixture of antibody-bound beads and egg extract was rotated on a wheel. Immunodepleted extracts were aliquoted, flash-frozen and kept at −80°C until use. One freeze-thaw cycle was found to have no effect on the extract’s TMEJ efficiency. Depletion efficiency was assessed by western blotting.

Polyclonal antibodies against *Xenopus* Mre11 were raised against full-length Mre11. Recombinant xMre11 was overexpressed in E. coli, purified as inclusion bodies and injected into rabbits following standard procedures (Pocono Rabbit Farm). Antibodies were affinity-purified from serum using the recombinant Mre11 coupled to AminoLink Coupling Resin (Thermo Fisher Scientific) according to manufacturer’s protocol and used for western blotting and immunodepletions.

### Immunoblotting and Phos-tag gel

All protein samples were boiled for 2 min at 95°C in Laemmli buffer (50 mM Tris-HCl pH 6.8, 2% SDS, 10% glycerol, 0.1% bromophenol blue, 5% β-mercaptoethanol). Samples were run on 4-15% Mini-PROTEAN precast gels (Bio-Rad) using Tris Glycine-SDS Running Buffer (25 mM Tris-HCl pH 8.3, 192 mM glycine, 0.1 % SDS).

To resolve phosphorylated Pol θ in Fig. 3C, PhosTag (Wako #AAL-107) SDS-PAGE gels were prepared with minor modifications [75]. Gels were composed of 3% acrylamide:bisacrylamide (37.5:1), 0.5% agarose, 200 mM Tris pH 7.0, 0.2% TEMED, 50 μM ZnCl2, 0.06% APS, 2.5 μM Phos-tag acrylamide. Running buffer contained 50 mM Tris, 50 mM Tricine, 0.1% SDS and 5 mM sodium bisulfate. Gels were transferred to 0.45 μm PVDF membranes (Thermo Scientific #88518) in transfer buffer (25 mM Tris pH 8.5, 192 mM glycine, 20% methanol, 0.1% SDS) at 500 mA for 1 h.

Membranes were blocked with 5% (w/v) non-fat milk in 1× PBST for 1h at room temperature, followed by incubation with primary antibodies diluted in 1× PBST (1:1000 for Pol θ, 1:10000 for Ku and Mre11) containing 1% BSA overnight at 4°C. Membranes were washed three times for 5 min with 1× PBST and incubated with goat anti-rabbit IgG horseradish peroxidase-conjugated antibody (Jackson ImmunoResearch) diluted 1:20,000 in 5% nonfat milk in 1x PBST for 45 min at room temperature. Membranes were washed extensively in 1x PBST, incubated for ~1 min with HyGLO chemiluminescent HRP antibody detection reagent (Denville), and imaged using an Amersham Imager 600 (GE Healthcare).

### Single stranded DNA pull down

A 150nt single stranded DNA oligonucleotide (9) with a biotin moiety attached to the 5′ terminus (biotin-ssDNA) was purchased from Integrated DNA Technologies. For each condition, 4 μL of streptavidin-coated magnetic beads (Sigma, Dynabeads M-280) were conjugated to biotin-ssDNA and 5 μL of free streptavidin-coated magnetic beads were added during pull down to facilitate sample recovery.

Beads were washed twice in 2x Bead Wash Buffer (10 mM Tris, pH 7.4, 2 M NaCl, 1 mM EDTA), and subsequently resuspended in 1x Bead Wash Buffer with 8 nM of the biotin-ssDNA at room temperature for 30 min. The DNA-bound beads were then washed twice with 2x Bead Wash Buffer, twice in Egg Lysis Blocking Buffer (10 mM HEPES, pH 7.7, 50 mM KCl, 2.5 mM MgCl2, 250 mM sucrose, 0.25 mg/ml BSA and 0.02% Tween20) and resuspended in 1.7 μl of Egg Lysis Blocking Buffer for each condition.

Extract was supplemented with the 30x ATP regeneration mixture (65 mM ATP, 650 mM phosphocreatine, 160 ng/μL creatine phosphokinase) to a 1x final concentration and 16 ng/μl B–CDK1 or B–CDK1 buffer. Each DNA-bead sample was mixed with 6.4 μL of the extract and incubated for 30 min at room temperature under slow rotation. At the end of the incubation, 1 μL of the reaction mixture was collected directly in 1x Laemmli sample buffer as whole extract sample. 6 μL of the reaction mixture was diluted into 40 μL of ice-cold Egg Lysis blocking buffer containing free beads. The beads were then washed three times in 200 μl of ice-cold Egg Lysis Washing buffer (10 mM HEPES, pH 7.7, 50 mM KCl, 2.5 mM MgCl2, 0.25mg/ml BSA and 0.03% Tween20) and resuspended in 1x Laemmli sample buffer as pull down samples.

### Quantifications of joining efficiency

All quantified conditions were performed three or more times. Gel images were quantified using Multi Gauge V3.0 and ImageJ. First, the intensity values for the larger repaired DNA products (I_joined_) and the remaining substrate (I_substrate_) were quantified. Next, the joining efficiency was calculated as the ratio of (I_joined_) to (I_joined_ + I_substrate_) at each time point. The percentage of joined DNA at 0 min (background) was subtracted from the joining efficiency at each subsequent time point to obtain the final absolute joining efficiency. The values were plotted with error bars representing the standard deviation from the mean.

---

**Figure S1.**
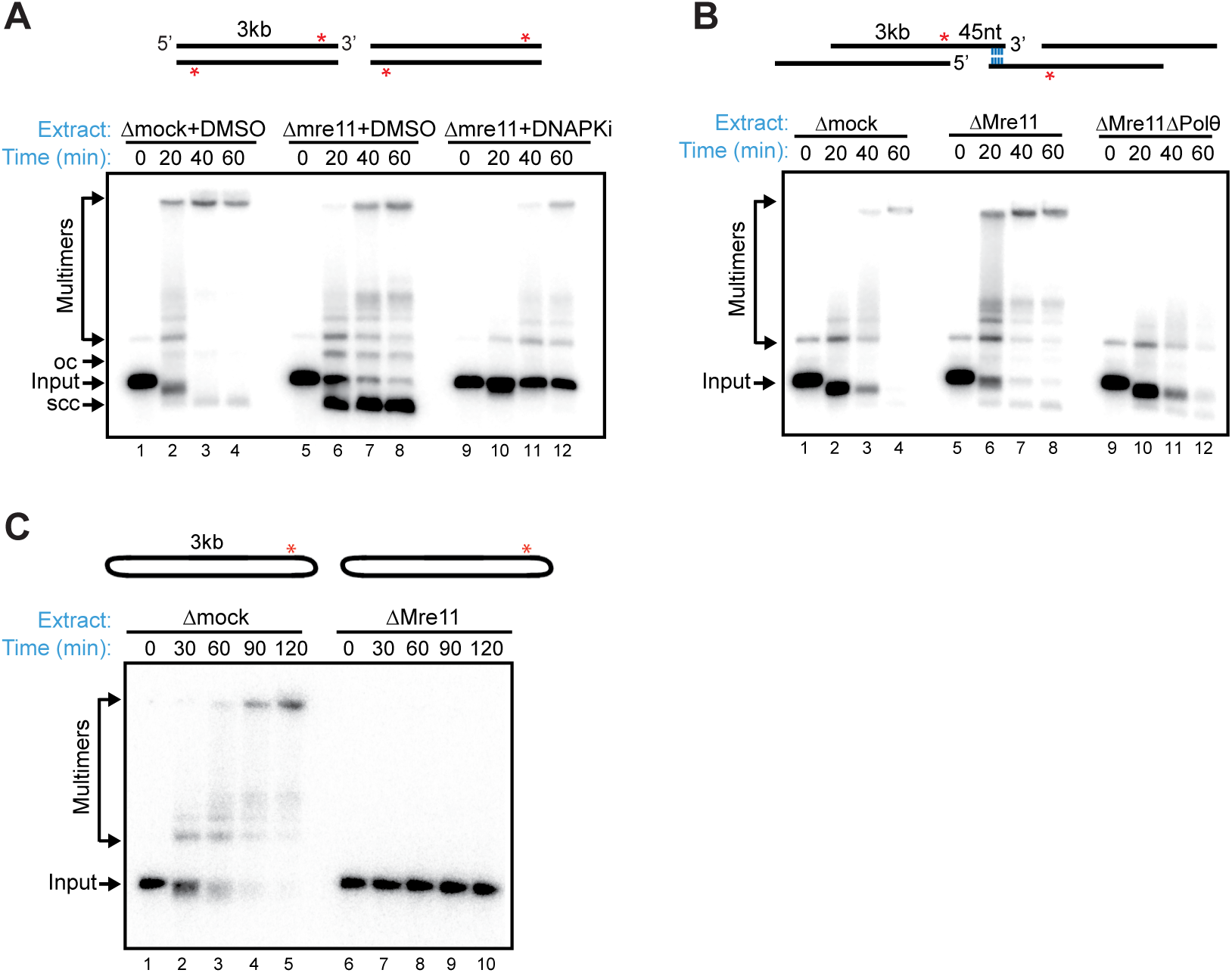
DSB repair pathway choice in *Xenopus* egg extract is dictated by DNA end structures. **(A)** Linear substrate was incubated with the indicated extracts (optionally supplemented with 10 μM DNA-PK inhibitor NU-7441), separated on a native agarose gel, and subjected to autoradiography. oc, open-circular products; scc, supercoiled closed-circular products. Red star, radiolabel. **(B)** Joining of a linear substrate with 45-nucleotides (nt) 3^’^ overhangs at both ends in the indicated extracts. Red star, radiolabel. Blue line, terminal microhomology (CCGG-3’). **(C)** Joining of a hairpin substrate in the indicated extracts. Red star, radiolabel.

**Figure S2.**
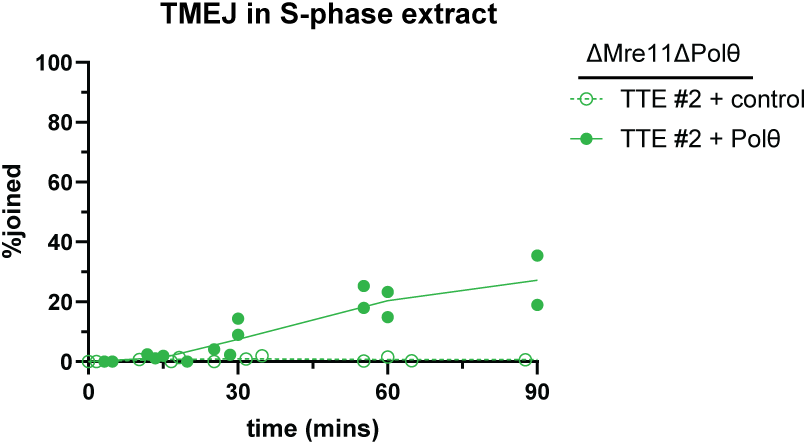
Rescue of TMEJ by recombinant Pol θ in interphase *Xenopus* egg extract. Same as Figure 1E, but using a different lot (#2) of TTE-expressed Pol θ. N=4 experiments. Note that in this case, maximum joining efficiency is ~20%.

**Figure S3.**
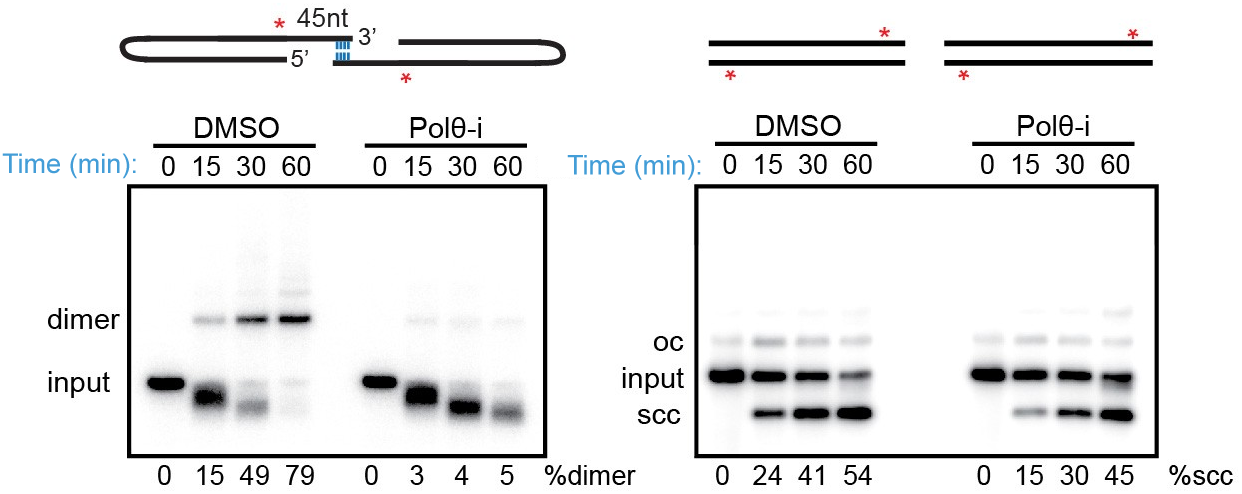
Suppression of TMEJ by a specific Pol θ Inhibitor. Joining of the 45OH (left) and the blunt-ended (right) substrates in the presence of DMSO or 10 µM Pol θ inhibitor ART558. oc, open-circular products; scc, supercoiled closed-circular products.

**Figure S4.**
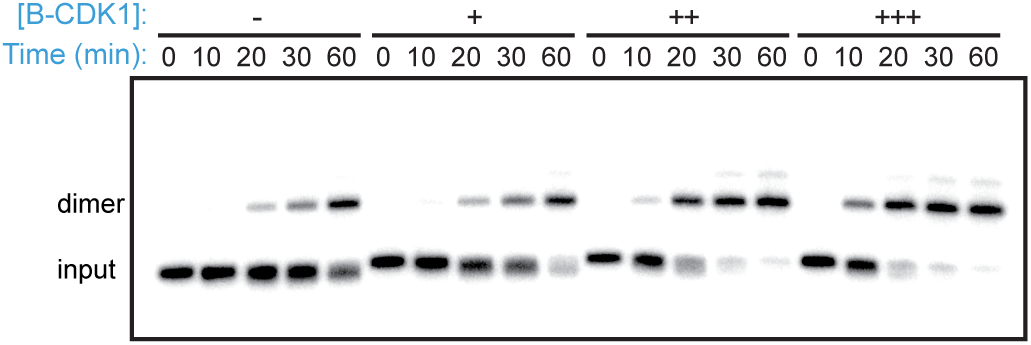
Cyclin B–CDK1 accelerates TMEJ. Non-denaturing gels showing time courses of joining of the 45OH substrate in Mre11-depleted NPE supplemented with increasing amounts of Cyclin B–CDK1 (0, 3, 8.3 and 25 ng/μl).

**Figure S5.**
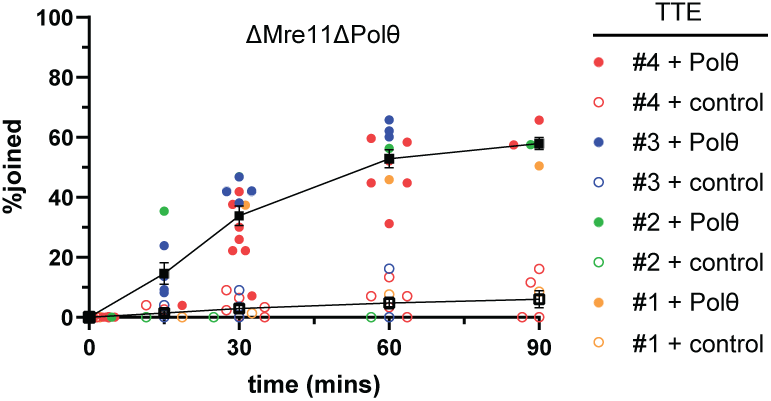
Rescue of TMEJ by recombinant Pol θ in mitotic *Xenopus* egg extract. Quantification of 45OH joining efficiency in NPE depleted of both Mre11 and Pol θ and supplemented with 16 ng/μl Cyclin B–CDK1 and wild-type Pol θ expressed using four different lots of tobacco-cell extract, including those used in Figures 1E and S2. N=13 experiments. Mean ± SEM.

**Table S1:**
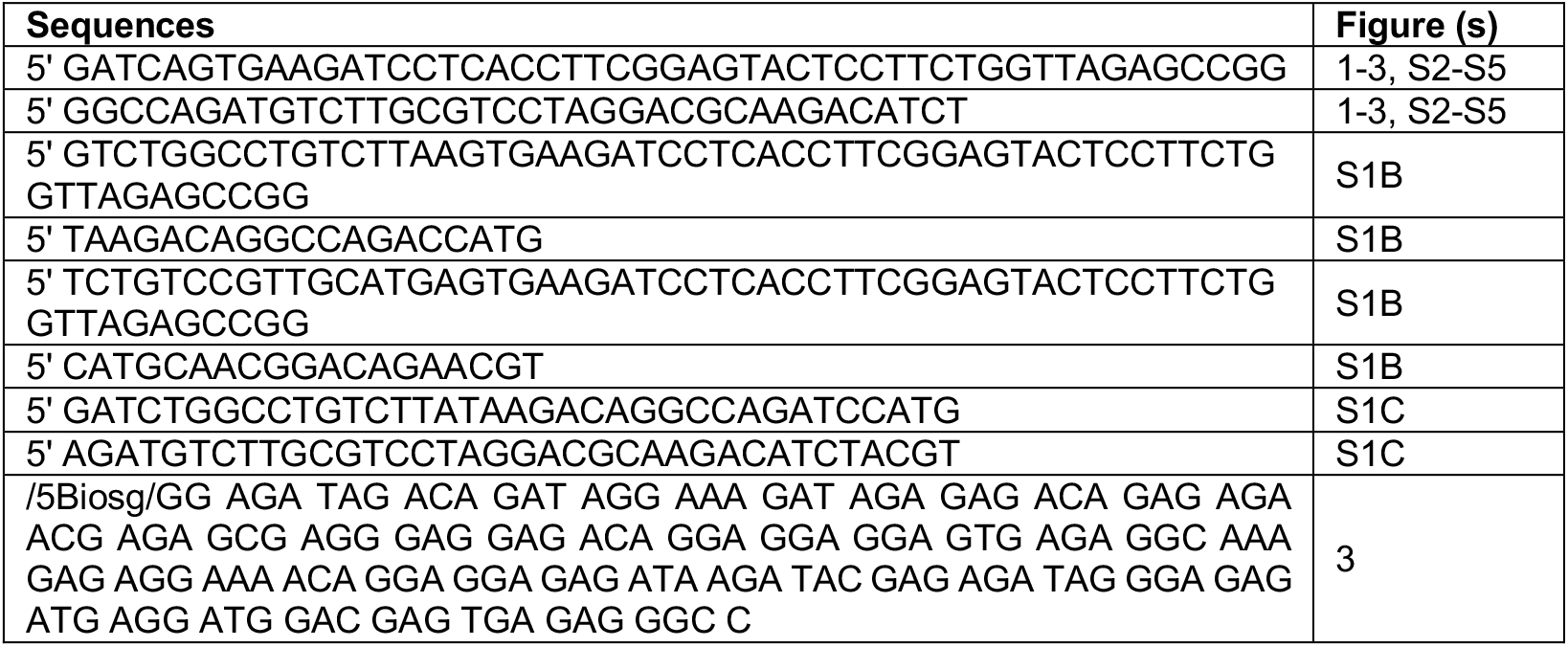
Oligonucleotides used to generate DNA substrates.

